# Trade-off between stimulation focality and the number of coils in multi-locus transcranial magnetic stimulation

**DOI:** 10.1101/2021.08.24.457503

**Authors:** Samuel Nurmi, Jere Karttunen, Victor H. Souza, Risto J. Ilmoniemi, Jaakko O. Nieminen

**Affiliations:** Department of Neuroscience and Biomedical Engineering, Aalto University School of Science, Espoo, Finland

**Keywords:** Transcranial magnetic stimulation, mTMS, multi-locus, transducer, coil, electric field, focality

## Abstract

**Objective:** Coils designed for transcranial magnetic stimulation (TMS) must incorporate trade-offs between the required electrical power or energy, focality and depth penetration of the induced electric field (E-field), coil size, and mechanical properties of the coil, as all of them cannot be optimally met at the same time. In multi-locus TMS (mTMS), a transducer consisting of several coils allows electronically targeted stimulation of the cortex without physically moving a coil. In this study, we aimed to investigate the relationship between the number of coils in an mTMS transducer, the focality of the induced E-field, and the extent of the cortical region within which the location and orientation of the maximum of the induced E-field can be controlled.

**Approach:** We applied convex optimization to design planar and spherically curved mTMS transducers of different E-field focalities and analyzed their properties. We characterized the trade-off between the focality of the induced E-field and the extent of the cortical region that can be stimulated with an mTMS transducer with a given number of coils.

**Main results:** At the expense of the E-field focality, one can, with the same number of coils, design an mTMS transducer that can control the location and orientation of the peak of the induced E-field within a wider cortical region.

**Significance:** With E-fields of moderate focality, the problem of electronically targeted TMS becomes considerably easier compared with highly focal E-fields; this may speed up the development of mTMS and the emergence of new clinical and research applications.

## Introduction

Transcranial magnetic stimulation (TMS) is a non-invasive brain stimulation method for diagnostics, therapy, and research [1]. TMS is typically administered with a round [2] or figure-of-eight [3] coil. Deng et al. demonstrated that there exists a trade-off between the focality and penetration of the induced electric field (E-field) [4]. Coils with optimal trade-offs between the E-field focality, E-field penetration, and stimulation energy can be designed computationally [5]. The most focal and least focal E-fields require the highest magnetic energies in the coil [6]. There is also a trade-off between the coil size and mechanical integrity, with the smallest coils being most susceptible to breaking during stimulation [7,8].

In multi-locus TMS (mTMS), the stimulation is administered with a transducer consisting of several coils that allow controlling the induced E-field pattern electronically [9,10]. This enables one to stimulate neighboring cortical areas with millisecond-scale interstimulus intervals [11] and to implement closed-loop stimulation paradigms [12,13]. TMS applications in which multiple nodes of a cortical network are stimulated in rapid succession, e.g., to study causal interactions in functional networks [14] or to administer therapy [15], may benefit from the ability to target specific cortical sites relatively far apart. An ideal mTMS transducer would allow controlling the E-field within a wide cortical region but contain only a small number of coils, as each coil requires an electronic circuit to drive current through its windings. A large coil set becomes problematic also if windings need to be stacked on top of each other. In this case, the furthermost coils would require excessive current amplitudes due to their suboptimal coupling to the brain [6,9]. Previously described mTMS transducers are sufficient for controlling the stimulation within an approximately 30-mm-diameter region [9,10]; however, many applications would benefit from a larger targeting region.

We investigated how the required number of coils in an mTMS transducer depends on the focality of the induced E-field and the extent of the cortical region within which the location and orientation of the maximum of the induced E-field can be controlled. We aimed to find a practical mTMS transducer design that would allow targeted cortical stimulation within a relatively large region. We hypothesized that by reducing the focality of the E-field, one can, with a fixed number of coils, control the location of its peak and its orientation within a wider cortical region.

## Methods

We modeled the human cortex as a spherical 70-mm-radius surface (12,995 points on the surface) and evaluated planar and spherically curved transducer geometries (Fig. 1A). We selected these geometries because planar transducers are easy to manufacture and fit all head shapes, whereas the curved transducer geometry allowed us to address the theoretical limits for transducers designed to follow the individual scalp shape. In this study, the transducer geometry refers to the overall transducer shape defined by the surface on which the coil windings reside (Fig. 1A); the coil winding patterns on these surfaces will be designed by applying an optimization procedure. The windings of the planar transducer were assumed to lie on a 300-mm-diameter disc (3,740-node triangular mesh), with its center 20 mm above the cortex. The windings of the curved transducer were constrained to the upper half of a 90-mm-radius sphere (3,047 nodes) concentric with the spherical cortex model.

**Figure 1.**
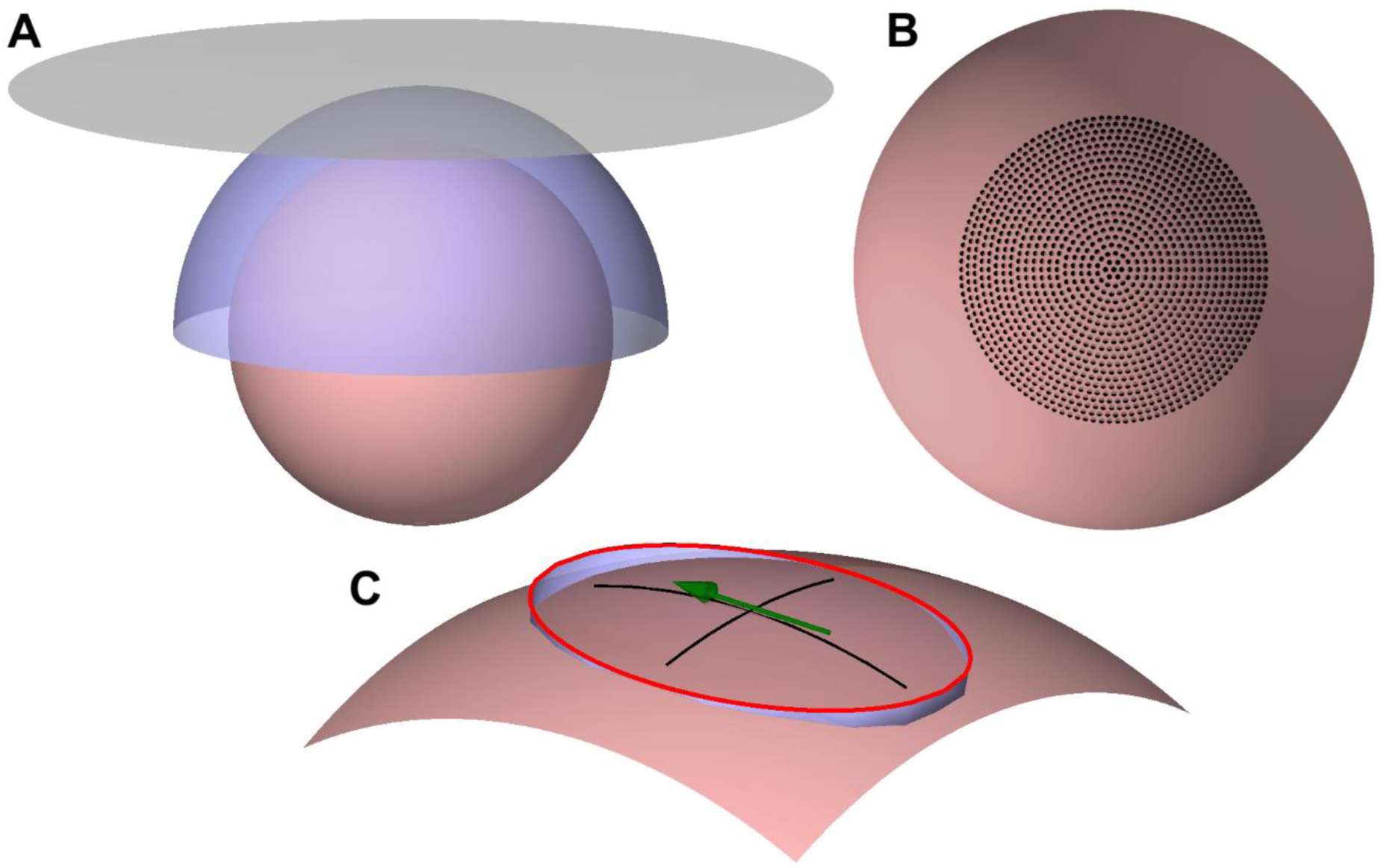
Optimization geometry. (**A**) The 70-mm-radius cortical surface (desaturated red), the spherical-cap coil surface (blue), and the planar coil surface (grey). (**B**) The target locations of the E-field maximum (black dots) on the cortical surface (desaturated red). (**C**) An illustration of the elliptical constraint region to control the E-field focality. An ellipse with a 3/2 aspect ratio (red) touches the cortex (desaturated red) at the end of its minor axes. The ellipse is projected to the cortex, with the projection path shown in blue. The green arrow illustrates the direction and location of the E-field maximum. The black lines are drawn along the projected minor and major axes of the ellipse.

With both transducer geometries, we aimed to have full control of the location and direction of the induced E-field maximum within a cortical region below the transducer. Thus, we defined a set of 1,276 target locations on the cortex (Fig. 1B). These targets were distributed on 20 concentric rings and at the center of the rings, which was on the axis of symmetry of the coil surfaces. The outermost ring had an 87-mm diameter measured along the cortical surface; on each ring, the distance between adjacent points matched the distance between the rings. At each target location, we considered only three E-field orientations (0°, 60°, and 120°), as a transducer capable of generating an E-field in all these directions can produce also intermediate E-field orientations by simultaneous activation of the coils.

To control the focality of the induced E-fields, we limited the extent of the region within which the E-field magnitude could exceed 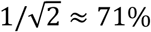 of the E-field magnitude at the target. To define this region, we first constructed an ellipse with a 3/2 ratio of its axes (Fig. 1C), as commercial coils produce E-field shapes close to this ratio [16] and minimum-energy coils tend to have E-field shapes with approximately a 3/2–2/1 aspect ratio [6]. We then placed the ellipse so that the points at the end of its minor axes were on the cortex, and finally projected the ellipse to the nearest points of the cortical surface (Fig. 1C). The direction of the longitudinal focality axis coincided with the direction of the E-field maximum. Outside the projected ellipse, the E-field magnitude was required to be less than 71% of the maximum. We investigated eight different E-field focalities by varying the width of the projected ellipse from 20 to 67 mm in 6.7-mm steps measured along the cortical surface. The diameter of the focal region along the direction of the E-field ranged from 30 to 100 mm in 10-mm steps. We refer to these focalities as Focality 20–30, Focality 27–40, …, and Focality 67–100, in which the two values refer to the width and length of the elliptical constraint region of a single stimulus, respectively.

For each of the 30,624 target E-field profiles (1,276 targets locations × 3 orientations × 8 focalities), we searched for the surface current densities on the planar and spherical-cap coil surfaces that produced the desired E-field pattern with the minimum energy by solving a convex optimization problem [6,17]. For each surface current density, we defined the energy as the magnetic field energy to induce a 100-V/m E-field at the target with a linear 100-µs current rise time. We extracted a subset of the solutions corresponding to the locations that were within a cortical region of a given diameter (i.e., a subset of the target rings shown in Fig. 1B). For each focality and both coil surfaces, the optimized current patterns were collected into a matrix whose columns represented the surface current density vectors. Then, each of these matrices was processed with the singular value decomposition (SVD) to obtain the left singular vectors describing possible coils of an mTMS transducer [9]. From each singular-vector set, we extracted the first vectors explaining 95% of the variance in the original coil set; their linear combinations would allow approximating any of the original optimized current density patterns [9]. Finally, coil winding paths were obtained as isolines of the optimized surface current density [18]. Note that a surface current density can be discretized into coils with different numbers of turns, the optimal number being dependent on many parameters, e.g., the electronics driving the coil and manufacturing constraints. In our illustrations, we have drawn the coils with such a number of turns that the winding patterns are easy to interpret.

## Results

### Single-coil optimization

Figure 2A shows the required stimulation energy of an optimized coil (or surface current density) as a function of the target location. With the spherical-cap coil, the stimulation energy for a given focality was found to be almost constant for all targets since the coil surface covered all targets similarly (i.e., all targets had the same coil–target distance and a wide amount of coil surface in all directions above them). With the planar coil, the required energy grew rapidly with an increasing distance between the target and the location directly below the transducer center. This growth in the stimulation energy was due to the increasing distance between the coil windings and the cortical target. As expected, for a given target, the required energy decreased when reducing the required E-field focality. There was, however, practically no difference between the five least focal E-field shapes with the spherical-cap coils, for which the curves in Fig. 2A overlap, or the Focality 60–90 and Focality 67–100 planar-coil E-fields, for which the curves in Fig. 2A follow each other closely. This is because with the least focal constraint regions the optimization was able to generate E-fields with the globally optimal focality; E-fields more focal or broader than this optimum would require more energy [6]. We also observe that all planar coils required more energy than the corresponding spherical coils, as theoretically shown by Koponen et al. [6]. The most focal E-fields studied (Focality 20–30) required over 200 J (or over 100%) more energy than the other E-field focalities, which would make such highly focal E-fields relatively difficult to produce in practice.

**Figure 2.**
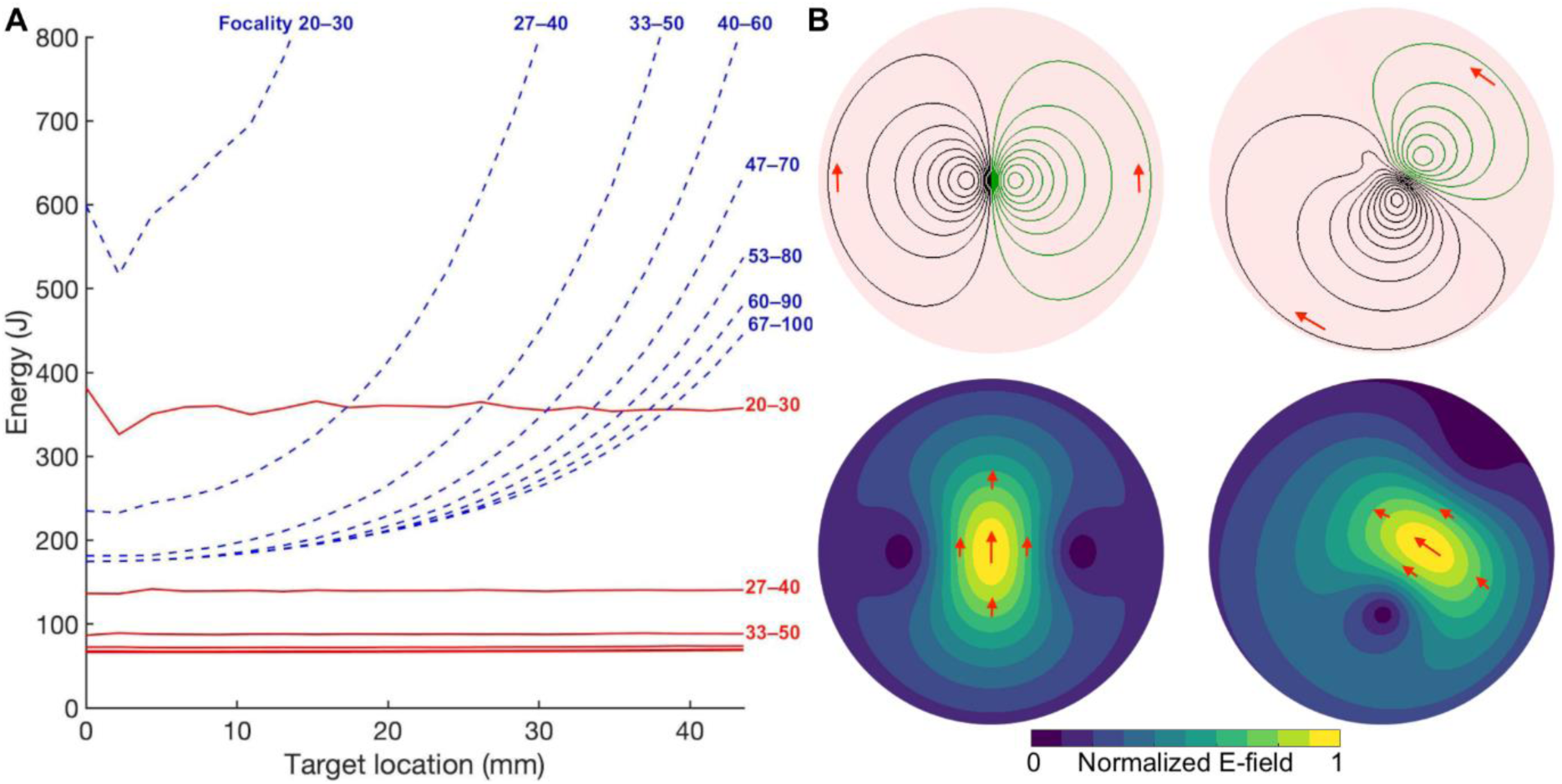
Single-coil optimization. (**A**) Magnetic field energy of the optimized single-coil surface current density patterns for planar (dashed blue lines) and spherical (solid red lines) coil geometries. The horizontal axis shows the distance along the cortical surface from the point below the transducer center to the target location. The annotations next to the curves indicate the focality they correspond to. The curves corresponding to the least focal E-fields in the spherical coil geometry are left unannotated. (**B**) Windings (top) in the planar coil geometry corresponding to the optimized E-field (bottom) with Focality 40–60 when the target is directly below the center of the coil surface (left) and 20 mm away from that location with the E-field rotated by 60° (right). The disk behind the windings illustrates the planar coil surface. Black and green windings represent clockwise and counterclockwise currents, respectively. The red arrows indicate the direction of the coil current and the induced E-field for a rising current.

As an example, Fig. 2B shows coil windings obtained as isolines of the optimized planar surface current density and the corresponding E-field (Focality 40–60) when the stimulation target was on top of the cortex and 20 mm away from that point (with a 60°-rotated E-field). For the stimulation of the topmost point, the energy optimization produced a symmetrical figure-of-eight coil as expected [17], whereas the translated target was optimally stimulated with an asymmetric coil.

### Optimized mTMS transducers

Figure 3 shows how the required number of coils in an optimized mTMS transducer increased as a function of the cortical target region and the explained variance. We observe, for example, that for Focality 40–60, five coils were sufficient to have 95% of the variance explained for a 43.5-mm-diameter target region for both transducer geometries (Figs. 3E–F). For Focality 40–60, the planar and spherical-cap transducers required 13 and 11 coils, respectively, to stimulate anywhere within the whole 87-mm-diameter target region (with the 95% variance threshold). Note, however, that for the planar geometry, the energy required to stimulate with that focality near the borders of the target region would be about fourfold compared to the energy required at the central region (Fig. 2). Figure 5I–J shows that for a transducer inducing less focal E-fields (Focality 67–100) the size of the available target region grew drastically. For example, with that focality, five coils would allow controlling the E-field maximum within cortical regions of 61 and 70 mm in diameter with planar and spherically shaped transducers, respectively (with the 95% variance threshold). Alternatively, the number of coils needed to control the E-field maximum within a given region could be reduced with less focal E-fields. From Fig. 3 we also notice that for a similar performance (E-field focality and control region size) a spherical-cap transducer required fewer coils than a planar transducer.

**Figure 3.**
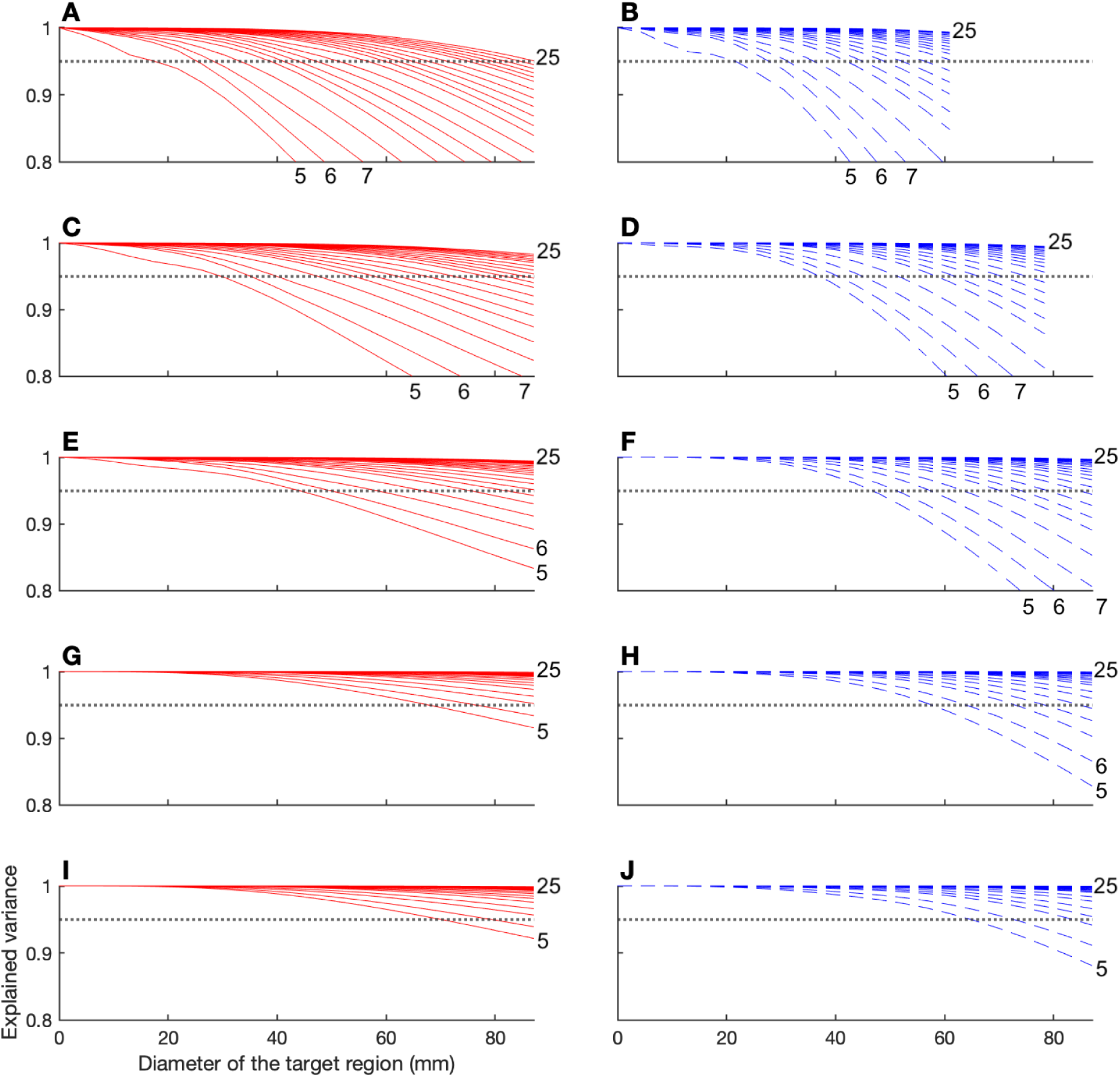
Number of coils needed in optimized mTMS transducers. The panels show data for spherical-cap (red) and planar (dashed blue) transducers with Focality 27–40 (**A, B**), Focality 33–50 (**C, D**), Focality 40–60 (**E, F**), Focality 53–80 (**G, H**), Focality 67–100 (**I, J**), respectively. The dotted grey line indicates the 95% explained variance threshold. The numbers next to the curves indicate the number of coils they correspond to. The data are plotted for transducers with 5–25 coils, as 25 coils allowed to reach the threshold of 95% of explained variance with all target-region sizes in all the visualized cases except for panel **B**. In panels **B** and **D**, the curves are cut at 61 mm and 83 mm, respectively, beyond which the required magnetic field energy exceeded 900 J.

Figure 4 shows examples of mTMS transducer windings and corresponding E-field distributions. In Fig. 4A, we show a planar 5-coil transducer with Focality 33–50 able to control the peak E-field within a 35-mm-diameter cortical target area (with 95% explained variance). This transducer has properties close to previously described 5-coil mTMS transducers [9,10]. In Fig. 4B, we show a planar transducer that can control the location and orientation of the peak E-field within a 61-mm-diameter cortical region (Focality 67–100). According to Fig. 2, at the borders of the cortical target region, this transducer would require only about 50% more energy than at a target below the transducer center.

**Figure 4.**
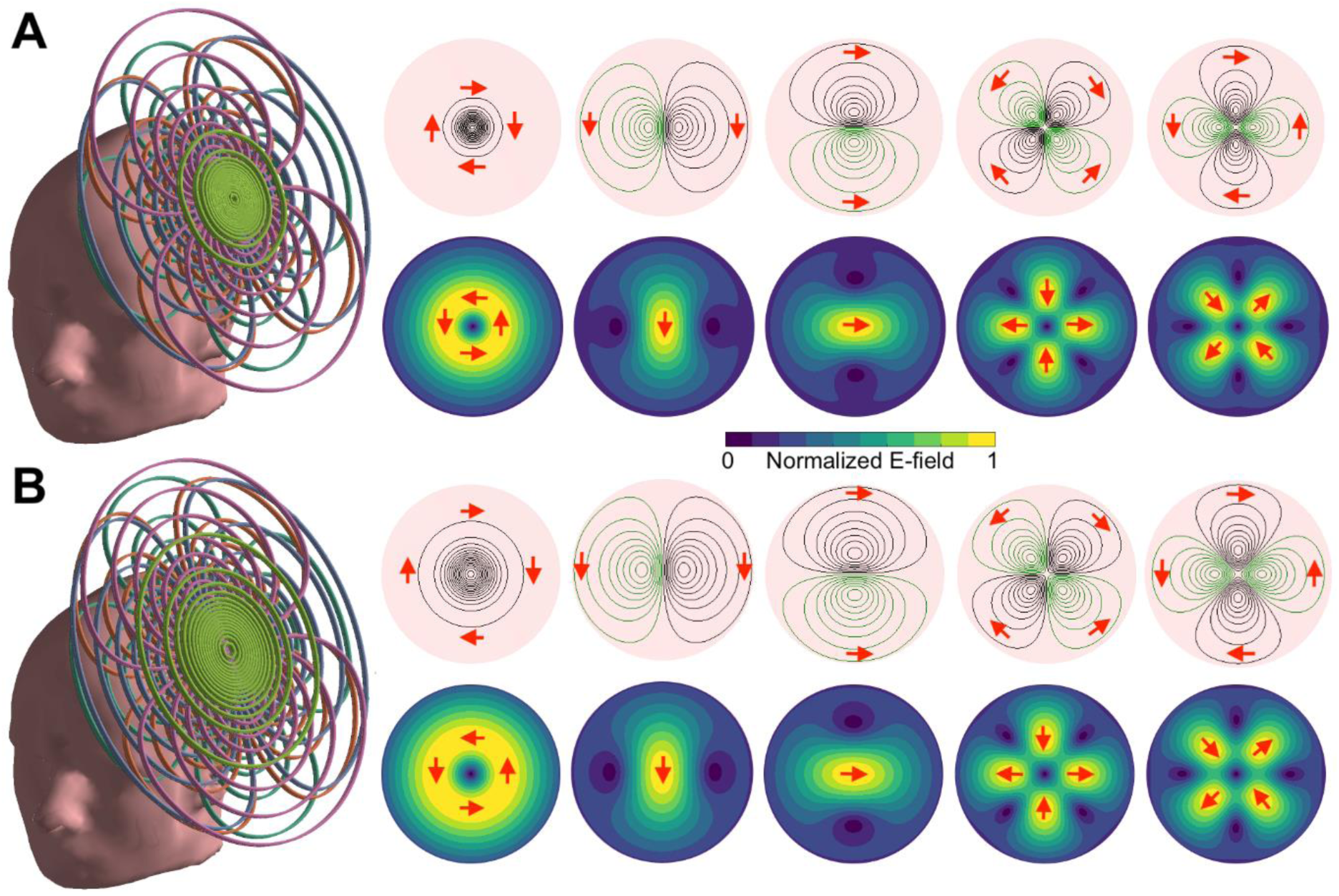
5-coil mTMS transducer designs. A planar 5-coil mTMS transducer for controlling the peak E-field within (**A**) a 35-mm-diameter cortical target region with Focality 33–50 E-fields and (**B**) a 61-mm-diameter cortical target region with Focality 67–100 E-fields. Left: The coil set placed tangential to the scalp. Right: The coil windings and the corresponding E-fields. Black and green windings represent clockwise and counterclockwise currents, respectively. The red arrows indicate the direction of coil current and the induced E-field for a rising current.

## Discussion

We analyzed the trade-off between the focality of the induced E-field and the extent of the cortical region that can be stimulated with an mTMS transducer consisting of a given number of coils. One may use this trade-off, e.g., to design a planar 5-coil mTMS transducer that can stimulate within a 60-mm-diameter cortical region. Such a region would be considerably larger than the cortical region available for previously described 5-coil mTMS transducers able to stimulate within a 30-mm-diameter region with more focal E-fields [9,10]. For a spherical-cap transducer able to control the stimulation target within an 87-mm-diameter region, the number of coils required dropped from 25 to 7 when the accepted E-field half-amplitude width was increased from 27 to 67 mm (Fig. 3). This is a substantial reduction in the requirements of the stimulator electronics and in the complexity of the coil windings. We foresee that for some applications, it may be useful to design a transducer for which the available E-field focality varies as a function of the target location. For example, one may envision an mTMS transducer that can induce highly focal E-fields in a central region and broader E-fields in the border regions. Such a transducer would allow implementing stimulation paradigms in which nearby targets in the central region are stimulated with highly focal E-fields providing the required level of specificity, while distant targets could be stimulated efficiently with broader E-fields.

The required level of E-field focality in TMS applications is uncertain. For example, presurgical cortical mapping [19,20] might benefit from highly focal E-fields. For other applications, e.g., treatment of major depression or stroke rehabilitation [21], it is more difficult to predict the best E-field focality. It may well be that future TMS researchers will have the option to select an mTMS transducers that best suits their needs, choosing between transducers capable of inducing the most focal E-fields within limited cortical regions and transducers with broader E-fields for larger cortical regions.

The E-fields resulting from the optimization were typically more focal than the elliptical constraint regions required. Mostly, the strongest (>71% of the maximum) E-field occurred within the whole length (i.e., along the major axis) of the elliptical constraint region, but in the perpendicular direction the distribution was narrower than the constraint, which allowed the optimization to reach the global minimum-energy solution. This was expected, given that E-field shapes with approximately a 3/2–2/1 aspect ratio require the lowest energies [6].

Spherical-cap transducers required less stimulation energy and could cover larger cortical areas than planar transducers. This was due to the matching shapes of the spherical cap and the cortex, which kept the coil–cortex distance small. With the planar-coil geometry, the increasing coil–cortex distance towards the border regions made it impossible to efficiently stimulate distant targets. This superiority of the spherical-cap geometry agrees with earlier theoretical results [6]. Planar transducers, however, are easy to manufacture and fit all head shapes.

The SVD processing of the single-coil surface current densities applied in this study produces coils that cover practically the whole available coil surface (see Fig. 4 for examples) [9]. As these coils would need to be stacked on top of each other, the maximum number of coils in such a transducer is limited in practice to less than about 10 coils, as otherwise the topmost coils would have very poor coupling to the brain. For mTMS transducers with a larger number of coils, one may apply, e.g., the varimax rotation [22] to obtain a coil set that could be built with a few layers, as suggested by Koponen et al. [9]. Such coil sets would contain partly overlapping near-round coils, with some characteristics in common to multi-coil TMS arrays suggested earlier [23–25].

Given that we analyzed the transducer properties only with simulations, our findings and the proposed transducers should be validated experimentally. In the present study, the transducers were designed with the help of a generic spherical head model, which generalizes over individual brain anatomies, simplifying the optimization [6,9,17,26]. However, when designing a curved transducer to be built, its geometry should preferably be based on the curvature of the scalp above the cortical target region to maximize the stimulation efficiency [6,17]. When an mTMS transducer is applied for brain stimulation, the E-fields are preferably calculated based on the individual cortical anatomy [10], regardless of the model applied in the transducer design, as E-field patterns in a realistic cortical geometry are rather complex [10,27].

## Conclusion

When designing an mTMS transducer, one should carefully evaluate and specify the desired level of E-field focality, as there is a trade-off between focality and the required number of coils needed to control the peak E-field within a given cortical region. Also, more focal E-fields require more energy than less focal ones. Ideally, mTMS transducers would also be designed to follow the scalp shape as closely as possible, as that results in benefits in the energy and the number of coils needed. In conclusion, with less focal E-fields, the problem of electronically targeted TMS eases considerably.

## Acknowledgements

This project has received funding from the Academy of Finland (Decisions No. 294625 and 327326), the European Research Council (ERC Synergy) under the European Union’s Horizon 2020 research and innovation programme (ConnectToBrain; grant agreement No. 810377), and the Jane and Aatos Erkko Foundation. We thank Science-IT at Aalto University School of Science for the computational resources.

## References

[1] Rossi S, Antal A, Bestmann S, Bikson M, Brewer C, Brockmöller J, Carpenter L L, Cincotta M, Chen R, Daskalakis J D, Di Lazzaro V, Fox M D, George M S, Gilbert D, Kimiskidis V K, Koch G, Ilmoniemi R J, Lefaucheur J P, Leocani L, Lisanby S H, Miniussi C, Padberg F, Pascual-Leone A, Paulus W, Peterchev A V, Quartarone A, Rotenberg A, Rothwell J, Rossini P M, Santarnecchi E, Shafi M M, Siebner H R, Ugawa Y, Wassermann E M, Zangen A, Ziemann U and Hallett M 2021 Safety and recommendations for TMS use in healthy subjects and patient populations, with updates on training, ethical and regulatory issues: Expert Guidelines Clin. Neurophysiol. 132 269–306

[2] Barker A T, Jalinous R and Freeston I L 1985 Non-invasive magnetic stimulation of human motor cortex Lancet 325 1106–7

[3] Ueno S, Tashiro T and Harada K 1988 Localized stimulation of neural tissues in the brain by means of a paired configuration of time-varying magnetic fields J. Appl. Phys. 10 5862–4

[4] Deng Z-D, Lisanby S H and Peterchev A V 2013 Electric field depth–focality tradeoff in transcranial magnetic stimulation: Simulation comparison of 50 coil designs Brain Stimul. 6 1– 13

[5] Gomez L J, Goetz S M and Peterchev A V 2018 Design of transcranial magnetic stimulation coils with optimal trade-off between depth, focality, and energy J. Neural Eng. 15 046033

[6] Koponen L M, Nieminen J O and Ilmoniemi R J 2015 Minimum-energy coils for transcranial magnetic stimulation: application to focal stimulation. Brain Stimul. 8 124–34

[7] Cohen D and Cuffin B N 1991 Developing a more focal magnetic stimulator. Part I: Some basic principles J. Clin. Neurophysiol. 8 102–11

[8] Yunokuchi K and Cohen D 1991 Developing a more focal magnetic stimulator. Part II: Fabricating coils and measuring induced current distributions J. Clin. Neurophysiol. 8 112–20

[9] Koponen L M, Nieminen J O and Ilmoniemi R J 2018 Multi-locus transcranial magnetic stimulation—theory and implementation Brain Stimul. 11 849–55

[10] Nieminen J O, Sinisalo H, Souza V H, Malmi M, Yuryev M, Tervo A E, Stenroos M, Milardovich D, Korhonen J, Koponen L M and Ilmoniemi R J 2021 Multi-locus transcranial magnetic stimulation system for electronically targeted brain stimulation bioRxiv 2021.09.20.461045

[11] Nieminen J O, Koponen L M, Souza V H, Stenroos M and Ilmoniemi R J 2019 Short-interval intracortical inhibition in human primary motor cortex: a multi-locus transcranial magnetic stimulation study Neuroimage 203 116194

[12] Tervo A E, Metsomaa J, Nieminen J O, Sarvas J and Ilmoniemi R J 2020 Automated search of stimulation targets with closed-loop transcranial magnetic stimulation Neuroimage 220 117082

[13] Tervo A E, Nieminen J O, Lioumis P, Metsomaa J, Souza V H, Sinisalo H, Stenroos M, Sarvas J and Ilmoniemi R J 2021 Closed-loop optimization of transcranial magnetic stimulation with electroencephalography feedback bioRxiv 2021.08.31.458148

[14] Groppa S, Werner-Petroll N, Münchau A, Deuschl G, Ruschworth M F S and Siebner H R 2012 A novel dual-site transcranial magnetic stimulation paradigm to probe fast facilitatory inputs from ipsilateral dorsal premotor cortex to primary motor cortex Neuroimage 62 500–9

[15] Arai N, Müller-Dahlhaus F, Murakami T, Bliem B, Lu M-K, Ugawa Y and Ziemann U 2011 State-dependent and timing-dependent bidirectional associative plasticity in the human SMA-M1 network J. Neurosci. 31 15376–83

[16] Nieminen J O, Koponen L M and Ilmoniemi R J 2015 Experimental characterization of the electric field distribution induced by TMS devices. Brain Stimul. 8 582–9

[17] Koponen L M, Nieminen J O, Mutanen T P, Stenroos M and Ilmoniemi R J 2017 Coil optimisation for transcranial magnetic stimulation in realistic head geometry Brain Stimul. 10 795–805

[18] Peeren G N 2003 Stream function approach for determining optimal surface currents J. Comput. Phys. 191 305–21

[19] Picht T, Schmidt S, Brandt S, Frey D, Hannula H, Neuvonen T, Karhu J, Vajkoczy P and Suess O 2011 Preoperative functional mapping for rolandic brain tumor surgery: comparison of navigated transcranial magnetic stimulation to direct cortical stimulation Neurosurgery 69 581–9

[20] Picht T, Krieg S M, Sollmann N, Rösler J, Niraula B, Neuvonen T, Savolainen P, Lioumis P, Mäkelä JP, Deletis V, Meyer B, Vajkoczy P and Ringel F 2013 A comparison of language mapping by preoperative navigated transcranial magnetic stimulation and direct cortical stimulation during awake surgery Neurosurgery 72 808–19

[21] Lefaucheur J P, Aleman A, Baeken C, Benninger D H, Brunelin J, Di Lazzaro V, Filipović SR, Grefkes C, Hasan A, Hummel F C, Jääskeläinen S K, Langguth B, Leocani L, Londero A, Nardone R, Nguyen J P, Nyffeler T, Oliveira-Maia A J, Oliviero A, Padberg F, Palm U, Paulus W, Poulet E, Quartarone A, Rachid F, Rektorová I, Rossi S, Sahlsten H, Schecklmann M, Szekely D and Ziemann U 2020 Evidence-based guidelines on the therapeutic use of repetitive transcranial magnetic stimulation (rTMS): An update (2014–2018) Clin. Neurophysiol. 131 474–528

[22] Kaiser H F 1958 The varimax criterion for analytic rotation in factor analysis Psychometrika 23 187–200

[23] Ruohonen J and Ilmoniemi R J 1998 Focusing and targeting of magnetic brain stimulation using multiple coils Med. Biol. Eng. Comput. 36 297–301

[24] Ruohonen J, Ravazzani P, Grandori F and Ilmoniemi R J 1999 Theory of multichannel magnetic stimulation: Toward functional neuromuscular rehabilitation IEEE Trans. Biomed. Eng. 46 646– 51

[25] Han B H, Chun I K, Lee S C and Lee S Y 2004 Multichannel magnetic stimulation system design considering mutual couplings among the stimulation coils IEEE Trans. Biomed. Eng. 51 812–7

[26] Souza V H, Nieminen J O, Tugin S, Koponen L, Baffa O and Ilmoniemi R 2021 Fine multi-coil electronic control of transcranial magnetic stimulation: effects of stimulus orientation and intensity bioRxiv 2021.08.20.457096

[27] Stenroos M and Koponen L M 2019 Real-time computation of the TMS-induced electric field in a realistic head model Neuroimage 203 116159

